# Gain-of-function Shh mutants activate Smo *in cis* independent of Ptch1/2 function

**DOI:** 10.1101/172429

**Authors:** Catalina Casillas, Henk Roelink

## Abstract

Sonic Hedgehog (Shh) signaling is characterized by strict non-cell autonomy; cells expressing Shh do not respond to their ligand. Here, we identify several Shh mutations that gain the ability to activate the Hedgehog (Hh) pathway *in cis*. This activation requires the extracellular cysteine rich domain of Smoothened, but is otherwise independent of Ptch1/2. Many of the identified mutations disrupt either a highly conserved catalytic motif found in peptidases or an a-helix domain frequently mutated in holoprosencephaly-causing *SHH* alleles. The expression of gain-of-function mutants often results in the accumulation of unprocessed Shh pro-peptide, a form of Shh we demonstrate is sufficient to activate the Hh response cell-autonomously. Our results demonstrate that Shh is capable of activating the Hh pathway via Smo independently of Ptch1/2, and that it harbors an intrinsic mechanism that prevents cell-autonomous activation of the pathway to favor non-cell autonomous signaling.

## Introduction

Sonic Hedgehog (Shh) is a signaling molecule indispensable for vertebrate embryonic development and adult stem cell maintenance. Impaired regulation of the Hedgehog (Hh) pathway can cause of various birth defects and diseases, including Holoprosencephaly (HPE) and a several of tumor types (Bale, 2002). Shh is synthesized as a 45 kDa pro-peptide encompassing a signaling “Hedge” domain and an intein-like “Hog” domain at the C-terminal end. Previous work has shown that the “Hog” domain mediates an autocatalytic cleavage of Shh, resolved only by the addition of a cholesterol moiety on the C-terminus of the 19 kDa “Hedge” domain (ShhN_Chol_) (Lee et al., 1994). ShhN_Chol_ becomes further modified with the attachment of a palmitoyl group to its N-terminus (Buglino and Resh, 2008). Lipid modified ShhN_Chol_ is then secreted from expressing cells through a mechanism requiring Dispatched1 (Disp1), Scube2, and ADAM metalloproteases. The release of ShhN_Chol_ from cell membranes requires the removal of its lipid modifications, which results in soluble and biologically active ligands capable of non-cell autonomous signaling (Jakobs et al., 2014; Ohlig et al., 2012).

In the absence of Shh ligand, the receptors Patched1 (Ptch1) and Patched2 (Ptch2) inhibit the signal transducer Smoothened (Smo) through a sub-stoichiometric mechanism (Alfaro et al., 2014a; Taipale et al., 2002; J. Zhang et al., 2001). Ptch1 and Ptch2, as well as Boc, Cdo, and Gas1, are required extracellular Shh receptors that regulate the Hh responses *in vivo* (Allen et al., 2011; Goodrich et al., 1997; Izzi et al., 2011). The binding interface between Shh and Ptch1, as well as the Shh antagonist Hhip (Bosanac et al., 2009), encompasses a Zn^2+^ ion coordinated within Shh that is part of a larger and highly conserved motif resembling a zinc peptidase catalytic domain (Hall et al., 1995a). The conserved catalytic site, and consequently any putative enzymatic activity, has been shown not to be required for non-cell autonomous, or *in trans*, signaling (Fuse et al., 1999); thus it is often referred to as a “pseudo active” site in Shh. The binding interface between Shh and its co-receptors, Boc, Cdo, and Gas-1, incorporates a portion of the catalytic domain that chelates two Ca^2+^ ions (McLellan et al., 2008). Accordingly, mutations that disrupt Ca^2+^ chelation (such as E90A (Izzi et al., 2011)) interfere with the binding of Shh to Boc, Cdo, and Gas1, and have been used to demonstrate the requirement for co-receptor binding to initiate the Hh response *in trans*.

The binding of Shh to a receptor complex on the surface of receptive cells leads to its internalization and the disinhibition of Smo, an event that results in changes to the subcellular distribution and activity of the signal transducer. The mechanism that links the extracellular binding of Shh to the activation of predominantly intracellular Smo is enigmatic (Aanstad et al., 2009; Bijlsma et al., 2012; Incardona et al., 2002; 2000; Milenkovic et al., 2009). Smo is a Class F G-protein coupled receptor (GPCR) and belongs to a receptor superfamily predominantly defined by Frizzleds (Frz), the canonical receptors of the Wnt pathway (Bhanot et al., 1996; Kristiansen, 2004). Smo and Frzs share over 25% sequence identity; both contain a characteristic heptahelical domain as well as a conserved extracellular Cysteine Rich Domain (CRD). Wnts bind to the CRD of Frz receptors through two distinct binding sites, one of which is a protein-lipid interface, to initiates signal transduction (Janda et al., 2012). Although Ptch1-mediated inhibition of Smo is thought to target a specific hydrophobic binding pocket within the heptahelical domain of Smo, the CRD of Smo can also regulate Hh pathway activity by binding to a variety of sterols- most of which cause an activation of the response (Myers et al., 2013; Nachtergaele et al., 2013; Nedelcu et al., 2013) (Corcoran and Scott, 2006; Huang et al., 2016; Xiao et al., 2017). The regulatory interactions involving the CRD of Smo demonstrate that activation of the Hh pathway is multifaceted, and that Ptch1-mediated inhibition is only part of a more complex regulation of Smo activity.

In this study, we find that expression of several Shh mutants activates the transcriptional Hh response cell-autonomously in *Ptch1*^LacZ/LacZ^;*Ptch2^−/−^* cells. *Ptch1*^LacZ/LacZ^;*Ptch2^−/−^* cells also retain the ability to migrate towards a source of ShhN, a non-canonical Hh response mediated by Smo. We demonstrate that this Shh-mediated activation events occur independently of all canonical Shh receptors, but requires the CRD of Smo. Many of the Shh mutants that activate the Hh response exhibit defects in processing resulting in the perdurance of Shh pro-protein. This Shh pro-protein retains its activity when expressed in the developing neural tube. Our results demonstrate that many Shh mutants are capable of activating the Hh response via Smo independent of Ptch1/2 function. As the amount of Shh pre-protein, but ShhN_Chol_, correlates with the level of cell-autonomous Smo activation, it appears that auto-processing of Shh turns the active Shh pre-protein into the inactive ShhN_Chol_. This form of Shh can be re-activated by the removal of the cholesterol moiety.

## Results

### Shh can activate the Hh response cell-autonomously independent of the canonical receptors

Signaling by Shh is strictly non-cell autonomous (García-Zaragoza et al., 2012; Shaw et al., 2009; Yauch et al., 2008). In SHH-induced tumors, the non-SHH-expressing cells are sensitive to Smo inhibitors, whereas cells expressing Shh are not. This notion of non-autonomy in Shh signaling is further supported by the observation that in embryos, cells that express Shh (e.g. cells in the notochord and floor plate) do not express *Ptch1*, a gene that is universally upregulated in cells responding to Shh. As *Ptch1* is not expressed in cells expressing Shh, we set out to test if the absence of this receptor causes the inability of cells to respond to Shh. To assure lack of all Ptch activity, we used *Ptch1*^LacZ/LacZ^;*Ptch2^−/−^* cells. These Ptch1/2-deficient cells do not induce the transcriptional Hh response when exposed to exogenously supplied ShhN (Roberts et al., 2016), which also ensures that any Hh pathway activity we measure is strictly cell-autonomous.

To similarly assess the requirement of Shh (co-)receptors to activate the Hh response, we used Shh-E90A (a mutant unable to bind to Boc, Cdo, and Gas1(Allen et al., 2011; Izzi et al., 2011)) and Shh-H183A a mutant that breaks the Zn^2+^ triad, which interferes with Hhip and Ptch1 binding. Both *Shh* and its mutant forms effectively induced the transcriptional Hh response after transfection in *Ptch1*^LacZ/LacZ^;*Ptch2^−/−^* cells (Figure 1A). To unambiguously demonstrate that the Shh co-receptors are not required for *in cis* activation of the Hh response, we used *Ptch1*^LacZ/LacZ^;*Ptch2^−/−^;Boc^−/−^;Cdo^−/−^;Gas1^−/−^;Shh^−/−^* cells. Upon neuralization ES cells with this genotype have an activated Shh response (Figure 1C), consistent with their lack of Ptch1/2. We transfected fibroblasts derived from the *Ptch1*^LacZ/LacZ^;*Ptch2^−/−^;Boc^−/−^;Cdo^−/−^;Gas1^−/−^;Shh^−/−^* ES cells with *Shh, ShhN* and their -*E90A* and -*H183A* mutant derivatives. We found that also in these receptorless cells the Hh response could be induced by these froms of Shh (Figure 1D). The ability of Shh and ShhN to induce the Hh response was shared with Ihh and Dhh (Figure 1E). These results demonstrate that activation of the Hh response *in cis* can occur independent of canonical (co-)receptors.

**Figure 1.**
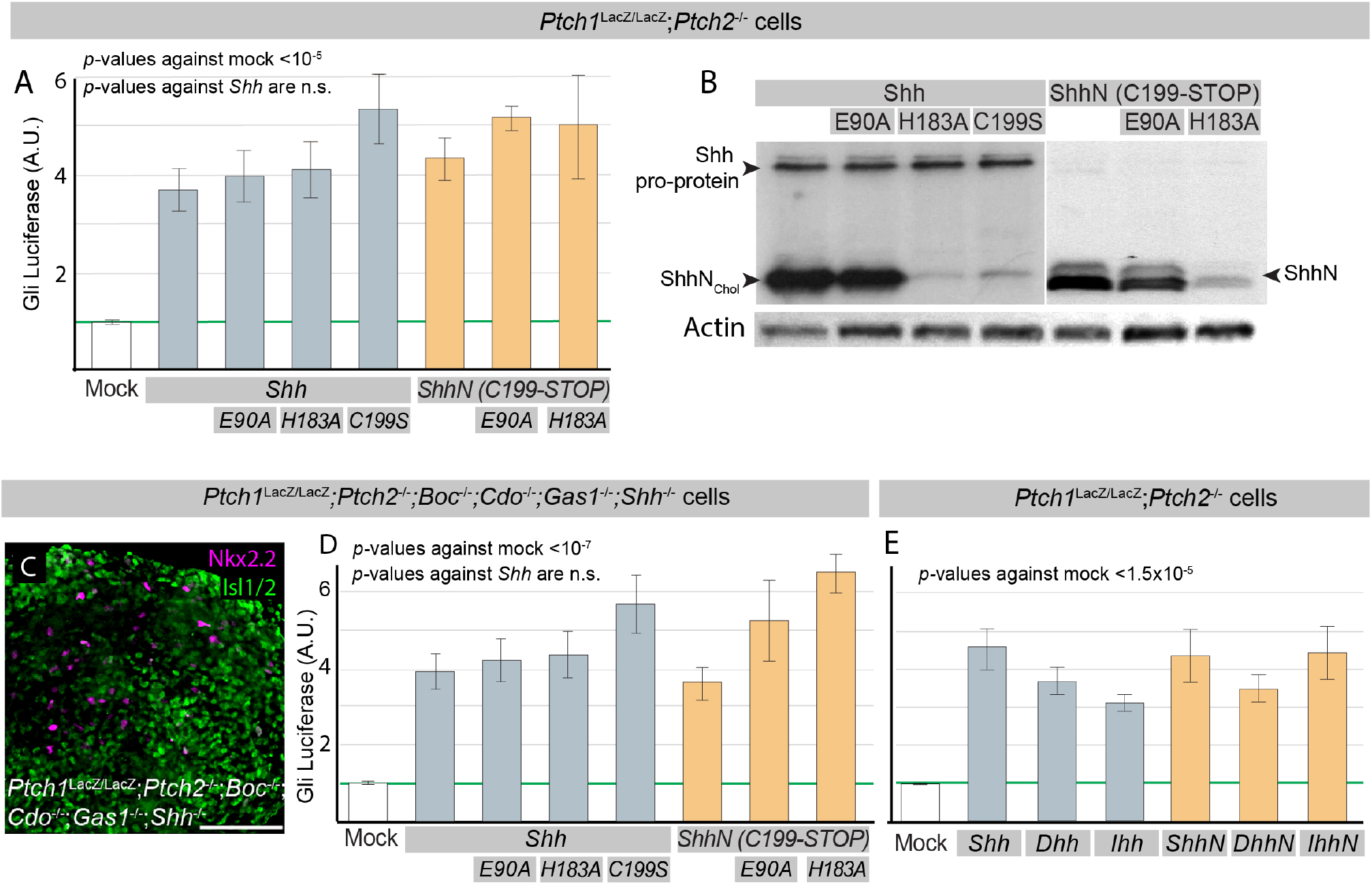
Shh mutants affecting the cholesterol moiety or the metal coordination sites gain the ability to activate the Hh response in cells lacking Hh receptors. A) *Ptch1*^LacZ/LacZ^; *Ptch2*^−/−^ cells were transfected with *Gli-Luciferase* (*Luc*) alone (Mock) or cotransfected with *Gli-Luc* and *Shh, Shh-E90A, Shh-H183A, ShhN, ShhN-E90A, or ShhN-H183A*. Luciferase levels in mock transfected *Ptch1*^LacZ/LacZ^;*Ptch2^−/−^* cells were set at “1”. (B) Western blot analysis of Shh, Shh-E90A, Shh-H183A, Shh-C199A, ShhN, ShhN-E90A, and ShhN-H183A protein expression in *Ptch1*^LacZ/LacZ^; *Ptch2^−/−^* cells, using an antibody directed against the N-terminal domain of Shh. (C) Neuralized Embryoid body derived from *Ptch1*^LacZ/LacZ^;*Ptch2^−/−^;Boc^−/−^;Cdo^−/−^;Gas1^−/−^;Shh^−/−^*mESCs. Expression of Nkx2.2 and Olig2 was assessed by immunofluorescence after 5 days in culture. (D) *Ptch1*^LacZ/LacZ^;*Ptch2^−/−^;Boc^−/−^;Cdo^−/−^;Gas1^−/−^;Shh^−/−^* cells were transfected with *Gli-Luc* alone (Mock) or co-transfected with *Gli-Luc* and *Shh, Shh-E90A, Shh-H183A, ShhN, ShhN-E90A*, or *ShhN-H183A*. Luciferase levels in Mock transfected cells were set at “1”. (I) *Ptch1*^LacZ/LacZ^; *Ptch2^−/−^* cells were transfected with *Gli-Luc* alone (Mock) or co-transfected with *Gli-Luc* and *ShhN, DhhN, IhhN, Shh, Dhh, or Ihh*. Luciferase levels in Mock transfected cells were set at “1”. All error bars are s.e.m., *p* values (Student t-test, 2 tailed) against mock are indicated were relevant, (A) n≥25, (D) n≥11, (E) n≥6, all independent biological replicates.

Shh protein was readily detected in transfected *Ptch1*^LacZ/LacZ^;*Ptch2^−/−^* cells (Figure 1B). In these cells, Shh is normally processed into a cholesterol-associated form (ShhN_Chol_) (19 kD), but full-length Shh pro-protein (45 kD) can be detected as well. Cells expressing Shh-H183A had little N-terminal processed Shh, and comparable levels of full length pro-protein (Figure 1B). It thus appears that the ability to induce the transcriptional Hh response *in cis* correlates with the presence of the full length form of Shh, and not with the N-terminal processed form. To determine if unprocessed Shh proprotein could mediate the activation of the Hh response *in cis*, we tested Shh-C199S, a form of Shh largely unable to undergo autoproteolysis (Figure 1B) and found that this mutant was an equally potent inducer of the Hh response (Figure 1A). As cholesterol-modified ShhN (ShhN_Chol_) is only activated upon shedding and removal of its lipid moieties (Ohlig et al., 2011), we tested a form of Shh that does not have the cholesterol moiety (Shh-C199* or ShhN), and does not require an activation event. We found that Shh-C199* retains the ability to induce the Hh response *in cis*. The mechanism by which Shh-E90A, -H183A and ShhN allow activation of the Hh pathway *in cis* appears to be related, as the double mutants (*ShhN-E90A* and *ShhN-H183A*) have the same activities as the single mutants (Figure 1A), and do not produce additive effects.

Together these results demonstrate that presumed loss-of-function mutations (Shh-H183A, Shh-E90A, Shh-C199S, ShhN-E90A, and ShhN-H183A), all have the ability to signal cell-autonomously in cells lacking Ptch1/2 receptors. Cell-autonomous activation of the Hh response by Shh does not require processing of the pro-protein, nor binding to its (co-)receptors.

### Shh mutants can activate the Hh response in the developing neural tube

To verify that Shh can activate the Hh pathway *in cis* independent of receptor function, we tested a variety of Shh mutants deficient in their ability interact with (co)-receptors in the developing neural tube. Along with Shh-E90A and Shh-H183A, we tested Shh-199A, a mutant form that cannot undergo autoproteolysis, and ShhN-C25S a mutant form lacking the N-terminal acyl group which has been shown to be unable to interact with Ptch1 (Tukachinsky et al., 2016). We assessed the activity of Shh mutants *in vivo* by co-electroporating *Shh, Shh-E90A, Shh-H183A*, or *Shh-199A*, together with *GFP* in developing chick neural tubes. Expression of Shh, Shh-E90A, and Shh-C199A caused an expansion of Nkx2.2 and Mnr2 domains, as well as repression of Pax7 expression (Figure 2, panels A-H), indicating that these forms of Shh are active, despite their inability to interact with canonical receptors. Electroporation of *Shh-H183A* does not result in activation of the pathway (Figure 2 C, G, K). However, we found that *ShhN-H183A* could induce the Hh response after electroporation (Figure 2 C’,G’,K’), thus indicating that correct Zn^2+^ coordination is not required to activate the Hh response *in vivo*. Consistent with the activity of *Shh-E90A*, we found that *ShhN-E90A* was active in this assay, as was *ShhN-C25S*. Altogether, the *in vivo* results support our findings *in vitro* demonstrating that the binding of Shh to Ptch1/2, Cdo, Boc or Gas1 is not necessary to activate the Hh response. They further demonstrate that Hh pathway activity is associated with the Shh pro-protein.

**Figure 2.**
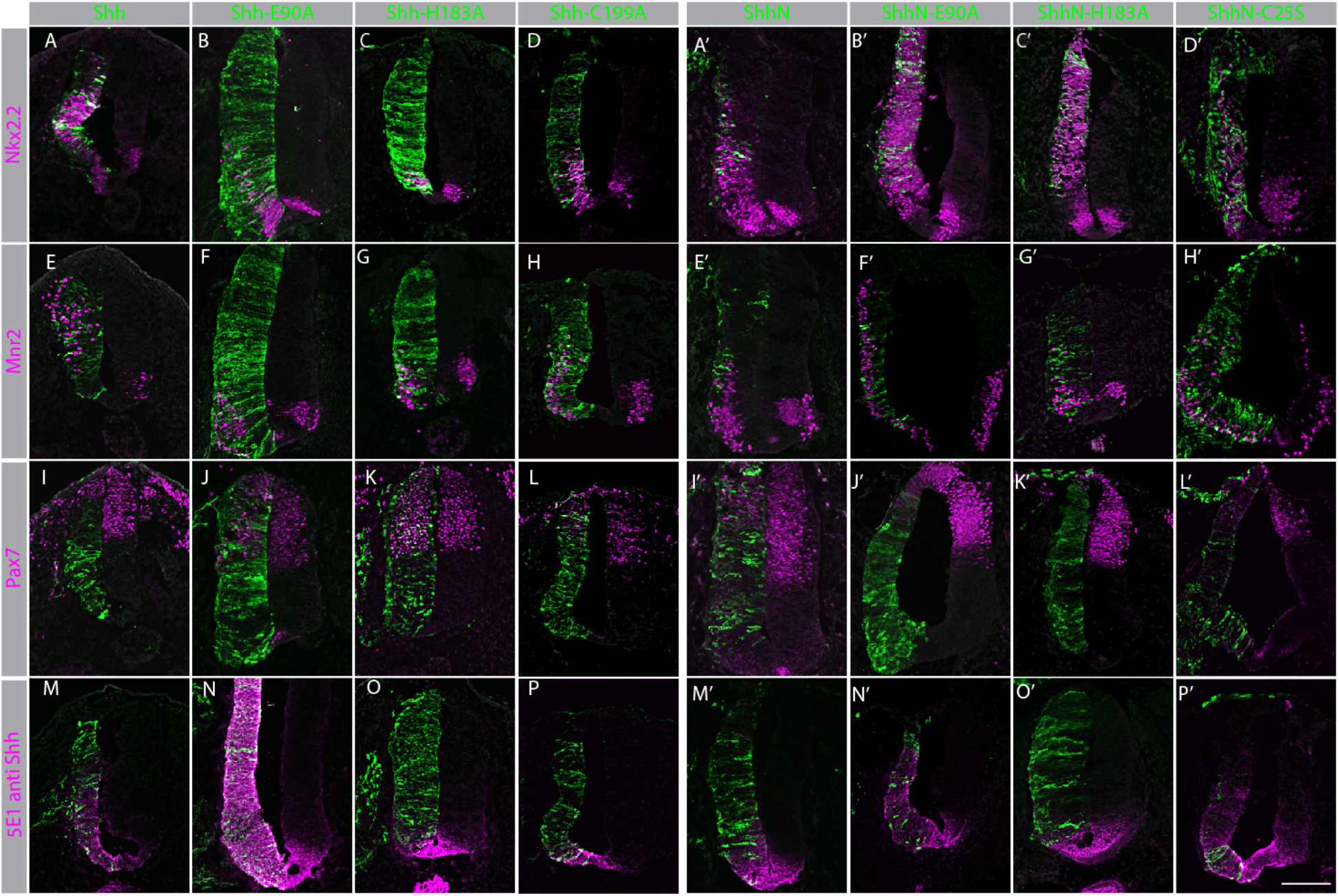
Shh and ShhN mutants activate the Hh pathway *in vivo* independent of receptor binding. (Panels A-P) Cross-sections of stage 20 HH chicken neural tubes co-electroporated with *GFP* and *Shh* (A, E, I, M), *Shh-E90A* (B, F, J, N), *Shh-H183A* (C, G, K, O), or *Shh-C199A (C199A* (D, H, L, P), labeled in green). Sections are stained with antibodies to Nkx2.2 (A-D), Mnr2 (E-H), Pax7 (I-L), and Shh (5E1) (M-P) labeled in magenta. (Panels A’-P’) Cross-sections of stage 20 HH chicken neural tubes co-electroporated at stage 10 with *GFP* and *ShhN* (A’,E’,I’,M’), *ShhN-E90A* (B’,F’,J’,N’), or *ShhN-H183A* (C’,G’,K’,O’). *ShhN-C25S* (D’,H’,L’,P’) labeled in green. Sections are stained with antibodies to Nkx2.2 (A-D, A’-D’), Mnr2 (E-H, E’-H’), Pax7 (I-L, I’-L’), and Shh (5E1) (M-P, M’-P’) labeled in magenta. Scale bar is 100μm.

The canonical model implicates the loss of Ptch1-mediated inhibition of Smo as the central mechanism by which the Hh response is activated. The observation that the Hh pathway can be activated by Shh in the absence of Ptch1/2 function raises the question of whether or not this pathway activation is mediated by Smo.

### *In cis* activation of the Hh response by ShhN requires Smo

The *Ptch1/2* null fibroblast line used in this study is derived from *Ptch1*^LacZ/LacZ^;*Ptch2^−/−^* mESCs (Roberts et al., 2016), which in turn are derived from *Ptch1*^LacZ/LacZ^ mESCs (Goodrich et al., 1997); consequently, these cells carry the *Ptch1:LacZ* null allele. As *Ptch1* itself is invariably induced in cells responding to Shh, the *Ptch1:LacZ* allele has found widespread use as a dependable readout of Hh pathway activation. To further assess the nature of cell-autonomous pathway activation, we also used *Ptch1*^LacZ/LacZ^ fibroblasts, used to study the of role of Ptch1 (Taipale et al., 2002), as a control. Transfection of *ShhN* into either cell line resulted in an activation of the transcriptional Hh response, i.e. a significant increase of transcriptional activity over the mock transfected cells. This activation could be blocked with the addition of vismodegib, a small molecule inhibitor of Smo (Sharpe et al., 2015) (Figure 3A). The ability of vismodegib to block Ptch1/2-independent activation of the Hh response, as a consequence of *ShhN* expression, demonstrates that this activity is mediated by Smo. This experiment was repeated using the *Gli-Luciferase* reporter as an independent readout in four independent cell lines of varying genotypes. The *Gli-Luciferase* reporter was invariably induced upon transfection with *ShhN*, regardless of whether the cells were Ptch1/2 proficient, *Ptch1* null, or lacked both Ptch1 and Ptch2 function (Figure 3B). The resulting Hh pathway activation could be lowered by vismodegib in all cell lines, further demonstrating the involvement of Smo in cell-autonomous signaling. *Gli-Luciferase* induction after ShhN expression confirms the results obtained using the *Ptch1:LacZ* allele for quantification (Figure 3A). As shown before, *Ptch1*^LacZ/LacZ^ fibroblasts have a moderate intrinsically upregulated Hh response after serum withdrawal (Taipale et al., 2002). This appears to be a unique characteristic of this particular cell line, as neither *Ptch1*^LacZ/LacZ^;*Ptch2^−/−^*fibroblasts, nor *Ptch1^−/−^;Disp1^−/−^;Shh^−/−^* fibroblasts exhibit this property (Figure 3 B,D).

**Figure 3.**
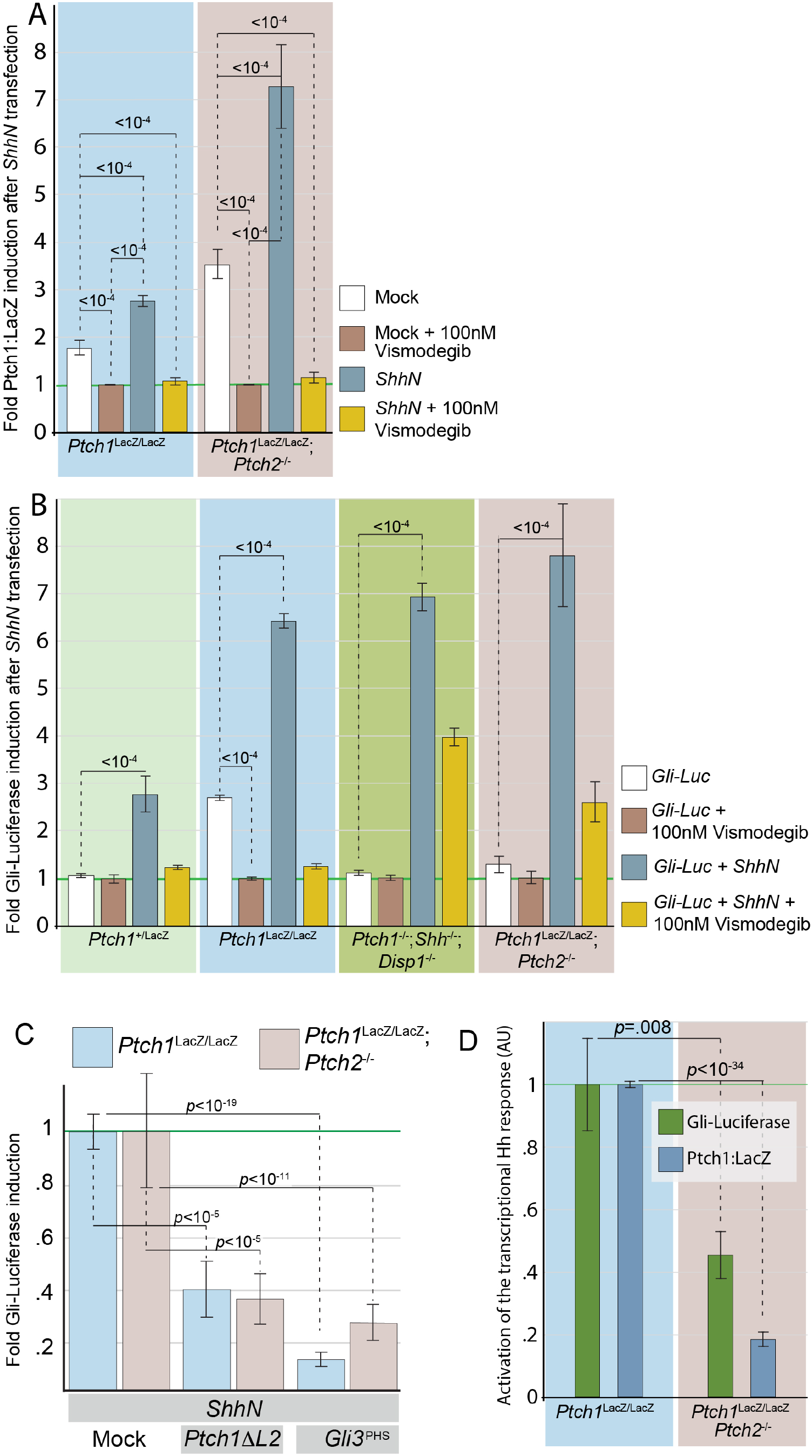
Activation of the Hh response by ShhN does not require Ptch1/2 activity. (A) *Ptch1*^LacZ/LacZ^ cells and *Ptch1*^LacZ/LacZ^;*Ptch2^−/−^* (indicated) were assessed for Ptch1:LacZ expression after mock transfection (white and brown bars) or *ShhN* transfection (blue and yellow bars). Each condition was treated with either a Smo inhibitor, 100 nM Vismodegib (brown and yellow bars), or a DMSO vehicle control (white and blue bars). β-Gal levels in mock transfected cells treated with 100 nM Vismodegib for each cell line were set at “1”. (B) *Ptch1*^+/LacZ^ cells, *Ptch1*^LacZ/LacZ^ cells, *Ptch1^−/−^;Shh^−/−^;Disp1^−/−^*, and *Ptch1*^LacZ/LacZ^;*Ptch2^−/−^*cells (indicated) were co-transfected with *Gli-Luc* and *GFP* (Mock; white/brown bars) or co-transfected with *Gli-Luc* and *ShhN* (blue and yellow bars). Each condition was treated with either a Smo inhibitor, 100 nM Vismodegib (brown and yellow bars), or a DMSO vehicle control (white and blue bars). Luciferase levels in mock transfected cells treated with 100 nM Vismodegib for each cell line were set at “1”. (C) Gli-luc levels were assayed in *Ptch1*^LacZ/LacZ^ (blue bars) and *Ptch1*^LacZ/LacZ^;*Ptch2^−/−^* (pink bars) transfected with *Ptch1ΔL2,or Gli3*^phs^ together with *Gli-luc* and *ShhN*. Luciferase levels in *ShhN*, mock co-transfected cells were set at “1” for each cell line. (D) Hh pathway activation in *Ptch1*^LacZ/LacZ^ and *Ptch1*^LacZ/LacZ^;*Ptch2^−/−^* was measured in parallel using *Gli-luc* and *Ptch1:LacZ*. Using both methods we find higher Hh pathway activation in the *Ptch1*^LacZ/LacZ^ cells than the *Ptch1*^LacZ/LacZ^;*Ptch2^−/−^* cells. Luciferase and LacZ levels in *Ptch1*^LacZ/LacZ^ cells were set at “1”. Error bars are s.e.m., p-values (Student t-test, 2 tailed) are indicated were relevant, n=6 for Gli-Luc assay, n≥25 for LacZ assay, all independent biological replicates. All error bars are s.e.m., *p*-values (Student t-test, 2 tailed) are indicated where relevant, (A) n≥4, (B) n≥9, (C) n≥13, n≥8 all independent biological replicates.

Cell-autonomous activation of the Hh response by *ShhN* in cells lacking Ptch1/2 activity could be inhibited by a dominant inhibitory form of Ptch1 and by the downstream transcription factor Gli3. Expression of *Ptch1ΔL2* (Briscoe et al., 2001) or *Gli3^PHS^* (Meyer and Roelink, 2003) prevented upregulation of the Hh response by ShhN in both *Ptch1*^LacZ/LacZ^ and *Ptch1*^LacZ/LacZ^;*Ptch2^−/−^* cells (Figure 3C). Ptch1 catalytically acts upon Smo (Taipale et al., 2002), which in turn acts via the Gli transcription factors (Ruiz i Altaba, 1998; Stamataki et al., 2005), thus these results indicate that ShhN expression activates Smo downstream of Ptch1/2 function and upstream of Gli function.

### The Cysteine Rich Domain of Smo is required for its activation of Shh

ShhN can act as a cellular chemoattractant (Angot et al., 2008; Bijlsma et al., 2007). The chemotactic response to Shh is directional and does not require *de novo* transcription or translation, nor does it require the function of Gli proteins (Bijlsma et al., 2007; 2008; Chinchilla et al., 2010; Lipinski et al., 2008). Chemotaxis towards Shh is mediated by Smo (Charron et al., 2003); however, it does not require the localization of Smo to the primary cilium, a prerequisite of the transcriptional response. Furthermore, chemotaxis can be mediated by forms of Smo unable to activate the transcriptional response to Shh (Bijlsma et al., 2012), indicating fundamental differences between these two activities of Smo. Translocation of Smo to the primary cilium involves Ptch1 function (Rohatgi et al., 2007), and we have previously shown that cells lacking *Ptch1* retain the ability to migrate towards ShhN (Alfaro et al., 2014b). Because Ptch1 and Ptch2 have overlapping functions (Alfaro et al., 2014b; Y. Lee et al., 2006; Roberts et al., 2016), it is not clear whether the chemotactic response in *Ptch1*^LacZ/LacZ^ cells is mediated by Ptch2 or if it is a Ptch1/2-independent signaling event. We tested the ability of *Ptch1*^LacZ/LacZ^;Ptch2^−/−^ fibroblasts to migrate towards a localized source of ShhN using a modified Boyden chamber (H.C. Chen, 2005). We found that *Ptch1*^LacZ/LacZ^;*Ptch2^−/−^* cells were indistinguishable from *Ptch1*^LacZ/LacZ^ cells in their ability to migrate towards ShhN (Figure 4A). Shh chemotaxis was largely abolished in *Ptch1*^LacZ/LacZ^;*Ptch2^−/−^;Smo^−/−^* cells; however, the ability to migrate towards FCS, our positive control, was retained (Figure 4A).

**Figure 4.**
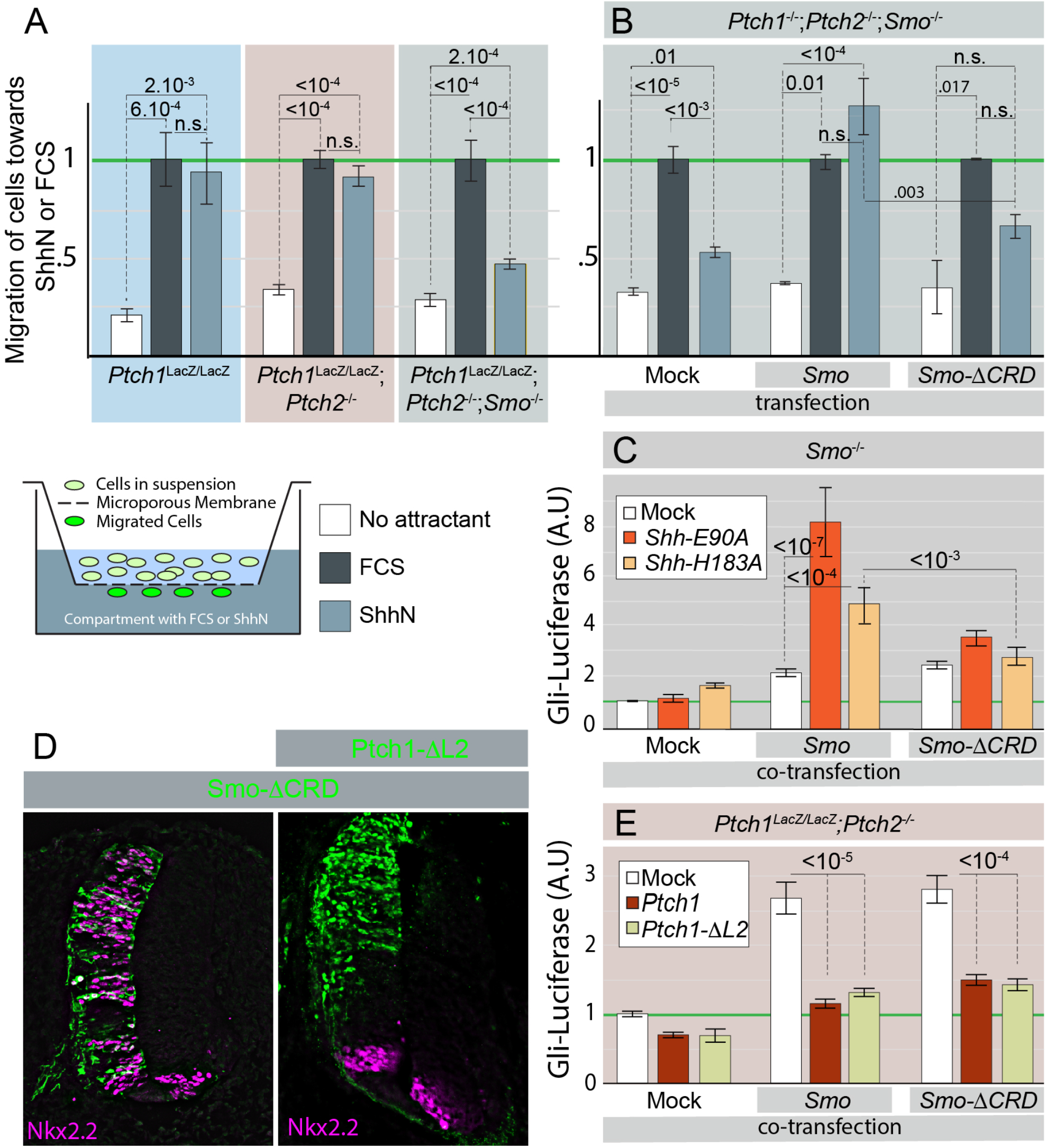
SmoΔCRD retains sensitivity to Ptch1-mediated inhibition in vitro and in vivo, but cannot be activated by Shh in aPtch1/2-independent manner. (A) Cells with the indicated genotypes were assayed in a modified Boyden chamber (diagram) for their ability to migrate towards FCS or ShhN. Migration was normalized to FCS for each condition. (B) *Ptch1^−/−^;Ptch2^−/−^;Smo^−/−^* cells were mock transfected, or transfected with *Smo* or *Smo-ΔCRD*. The transfected cells were assayed in a modified Boyden chamber for their ability to migrate towards FCS or ShhN. (C) *Smo^−/−^* cells were transfected with *Gli-Luc*, and co-transfected with *Smo* or *Smo-ΔCRD and Shh-E90A* or *Shh-H183A* as indicated. Gli-Luciferase levels were quantified, and the levels in double mock transfected cells set at “1”. (D) Stage 10 chicken embryos were electroporated with *Smo-ΔCRD* (green) or co-electroporated with *Smo-ΔCRD* and *Ptch1-ΔL2 (Green)*. The ventral marker NKX2.2 was visualized (purple). (E) *Ptch1*^LacZ/LacZ^;*Ptch2^−/−^* cells were transfected with *Gli-Luc*, and co-transfected with *Smo* or *Smo-ΔCRD, and Ptch1* or *Ptch1-ΔL2* as indicated. Gli-Luciferase levels were quantified, and the levels in *mock/mock/Gli-Luc* transfected cells set at “1”.

We could restore the ability of *Ptch1*^LacZ/LacZ^;*Ptch2^−/−^;Smo^−/−^* cells to migrate towards ShhN by transfecting them with wild type *Smo*, demonstrating the sufficiency of Smo to mediate Shh chemotaxis independent of Ptch1/2 function. (Figure 4A). However, transfection of a form of Smo lacking its CRD (*Smo-ΔCRD*) did not restore the chemotactic response in *Ptch1*^LacZ/LacZ^;*Ptch2^−/−^;Smo^−/−^* cells. These results demonstrate that Smo can recognize ShhN in the extracellular space to mediate the chemotactic response in the absence of Ptch1/2, and the CRD of Smo is required for this signaling event.

We used a *Smo^−/−^* fibroblast cell in lieu of the *Ptch1*^LacZ/LacZ^;*Ptch2^−/−^;Smo^−/−^* cell line to assess the requirement of the CRD of Smo to mediate the cell-autonomous transcriptional response. This was necessitated by our experience that *Ptch1*^LacZ/LacZ^;*Ptch2^−/−^;Smo^−/−^* cells do not survive the four days of serum deprivation necessary for the transcriptional assays. To ensure that we strictly measured cell-autonomous Ptch1/2-independent signaling, we used the *Shh-E90A* and *Shh-H183A* mutants that cannot signal non-cell autonomously. As expected, transfection of either *Shh-H183A* or *Shh-E90A* alone in *Smo^−/−^* cells did not result in the induction of the Hh response (Figure 4C). Transfection of *Smo* or *Smo-ΔCRD* alone caused a small upregulation of the Hh response. Consistent with earlier results (Figure 1A), we found that co-transfection of *Shh-E90A* or *Shh-H183A* together with *Smo* caused a strong induction of the Hh response *in cis*. In contrast, neither Shh-H183A nor Shh-E90A was capable of synergizing with Smo-ΔCRD to induce the Hh response, further demonstrating that the activation of Smo by Shh requires the CRD of Smo.

Despite its inability to respond to Shh, Smo-ΔCRD remains subject to inhibition by Ptch1. Mis-expression of *Smo-ΔCRD* in the developing chick neural tube after electroporation causes an upregulation of the Hh response (Figure 4D), consistent with earlier reports that Smo-ΔCRD retains a relatively high level of basal activity in vivo (Kwong et al., 2014). The activity of Smo-ΔCRD was suppressed by co-expression of *Ptch1ΔL2*, coding for a dominant-inhibitory form of Ptch1 (Briscoe et al., 2001) (Figure 4D). Similarly, expression of either *Smo* or *Smo-ΔCRD* in *Ptch1*^LacZ/LacZ^;*Ptch2^−/−^*cells results in transcriptional Hh activity, which can be inhibited by co-expression with *Ptch1* or *Ptch1ΔL2* (Figure 4E). Together these results that the CRD of Smo is required for its activation by ShhN, but not for its repression by Ptch1.

### The “pseudo-active” site of Shh has a role in Shh function other than receptor binding

Shh-E90 and -H183 are involved in the binding of Ca^2+^ and Zn^2+^ metal ions, respectively, and the coordination of the Zn^2+^ ion by H141, D148, and H183 (the “triad”, Figure 6A) resembles that of Zn^2+^ peptidases, (Hall et al., 1995a) including Thermolysin (Rebollido-Rios et al., 2014). Hhs belong to a larger family of DD-transpeptidases (van Heijenoort, 2011) that has many members in prokaryotes. In general, such carboxypeptidases resolve the acyl-enzyme intermediate with a nucleophile other than OH^−^, resulting in a sidechain addition (Fonzé et al., 1999). Peptidase activity of Shh is not required for activation of the Hh response (Fuse et al., 1999), (Figure 1 A, Figure 2), and the Zn2+ coordination was therefore often referred to as a “pseudo-active” site (Bosanac et al., 2009; Maun et al., 2010).

Figure 1B shows that Shh-H183A does not undergo autoproteolytic processing. To assess if the Zn^2+^ coordination triad is required for the formation of ShhN_Chol_ we also made Shh-H141A and Shh-D148A, and confirming earlier observations on SHH-H140P (Traiffort et al., 2004), we found that Shh-H141D did not auto-process, and D148A processed very poorly (Figure 5A). Together, this shows that correct Zn^2+^ coordination is necessary for processing the Shh pro-protein into ShhN_Chol_.

**Figure 5.**
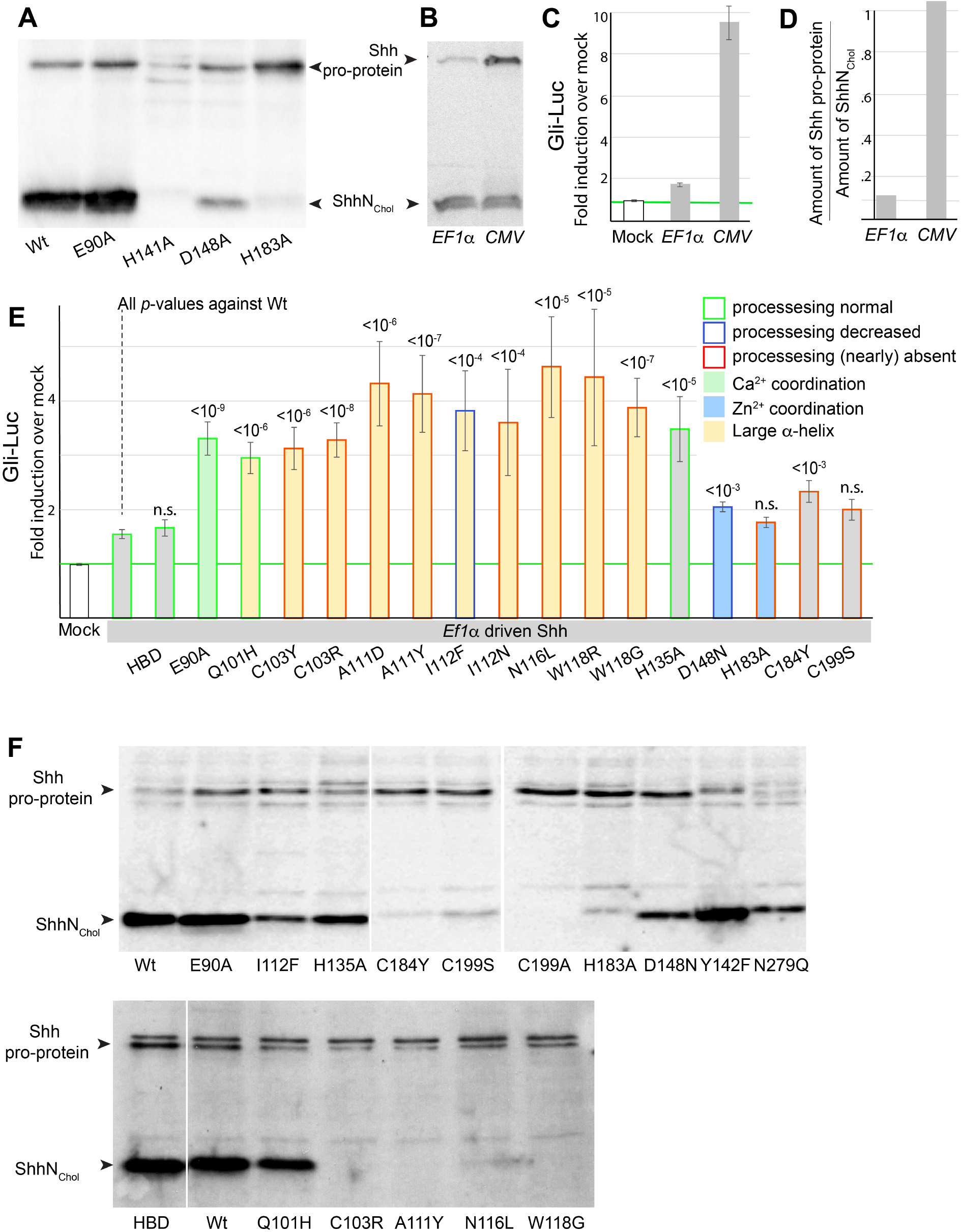
Shh processing is not required for the cell-autonomous activation of the Hh response. (A) Mutations in the Zn^2+^ coordination triad (H141A, D148A, H183A) severely compromise, or prevent Shh processing. E90A (affecting Ca^2+^ coordination) undergoes normal processing. (B) Shh expression levels (high for the *CMV* and low for the *EF1a* promoter) affect the level of Shh pro-protein more so than ShhN_Chol_. (C) The induction of the cell autonomous Hh response correlates with the amount of observed Shh pro-protein, but not Shh_Chol_. (D) Ratio of Shh pro-protein over ShhN_Chol_. (E) All Shh mutants are driven off the *EF1a* promoter. Activity to induce the Hh response *in cis* of several holoprosencephaly mutants located in the large a-helix (C103Y, C103R A111D, A111Y, I112F, I112N, N116L, W118R, W118G), mutants involved in Zn^2+^ coordination (D148N, H183A), E90A (Ca^2+^ coordination), H135 (near the catalytic domain), C184A (commonly mutated in holoprosencephaly), and C199S (cleavage site for auto-processing), Shh HBD (K38S, K39S, K179S) was used as a control. (F) Western blot analysis of EF1a-driven Shh mutants assessed in F. Note that the ability to auto-process is often impaired in a-helix and Zn^2+^ coordination mutants.

In Zn^2+^ peptidases, the E177 equivalent abstracts a proton from the catalytic water at the Zn^2+^ coordination domain, which is followed by a nucleophilic attack of the OH^−^ on the peptide backbone. Shh-E177A is, therefore, predicted to lack the intrinsic peptidase activity. Published analysis of this mutant has revealed two interesting properties. First, Shh-E177A is unable to mediate Shh signaling from the notochord to the overlying neural tube (*in trans*), but is more capable than wt Shh to induce the Hh response when expressed in the developing neural tube (likely *in cis*) (Himmelstein et al., 2017). Second, ShhN-E177A is more stable in solution than ShhN (Rebollido-Rios et al., 2014). We found that the Shh-E177A pro-protein was processed normally (not shown). Thus although the Zn^2+^ triad is required for processing of the Shh pro-protein, this does not appear to proceed via a canonical intrinsic peptidase activity. This is further supported by our finding that after co-expressing the obligatory pro-proteins Shh-C199A (unable to be cleaved) and Shh-H183A (unable to cleave) we could not detect any ShhN_Chol_ (not shown).

Both H141 and D148 are found mutated in holoprosencephaly, a congenital syndrome that can be caused by aberrant SHH signaling (Hehr et al., 2010; Odent et al., 1999; Roessler et al., 2009). It was observed before that several holoprosencephaly mutants prevent processing of the SHH proprotein (Traiffort et al., 2004), indicating that the perdurance of the SHH pro-protein might contribute to this congenital syndrome. As we found that the obligatory Shh pro-protein (Shh-C199A) was able to induce the Hh response in vivo we tested if failure of Shh processing could alter the Hh response. The autocatalytic processing of the Shh pro-protein is relatively slow, and to minimize potential overexpression artefacts we tested Shh expressed using the human *Elongation Factor 1α* (*EF1α*) promoter, which is much weaker than the *CMV* promoter used in previous experiments. When driven by the *EF1α* promoter, we found very little activity associated with Shh, unlike *Shh* driven by the *CMV* promoter (Figure 5C). Analysis of the protein expressed in these transfected cells (Figure 5B) revealed that the amount of observed Shh pro-protein correlates with the level of Hh pathway induction; this is in contrast to the amount of observed ShhN_Chol_, which was not correlated with pathway activity (Figure 5C,D). These observations further advance the notion that ShhN_Chol_ is unable to induce the Hh response *in cis* and that the Shh pro-protein can activate the Hh response.

The low level of Hh pathway induction observed in cells expressing *EF1α*-driven *Shh* allowed us to assess if Shh mutations result in a gain-of-function phenotype, as measured by activation of the transcriptional Hh response *in cis*. We tested mutations affecting the Zn^2+^ coordination domain, the Ca^2+^ coordination domain, mutations affecting the putative peptidase activity, and a series of holoprosencephaly mutants, in particular those in the large a-helix that dominates the face of Shh opposite to the Zn^2+^ coordination domain, and some mutations (W118R/G and C184F) tested (Traiffort et al., 2004). All these mutants were directly introduced into the *Shh* clone that had little activity after transfection. We found that Zn^2+^ coordination triad mutants (D148N, and H183A, blue bars, Figure 5E), caused no or minimal activation of the Hh response as compared to wild-type Shh. In contrast, E90A and H135A gained the ability to induce Hh response *in cis*, while being processed normally. Traiffort demonstrated that W118R/G does not process (Traiffort et al., 2004). Tryptophan118 is at the C-terminal end of the large a-helix. This helix is enriched in mutations found in holoprosencephaly patients (Figure 6C), and we tested Q101H, C103Y/R, A111D/Y I112F/N, N116L and W118R/G. All these of Shh point mutations are gain-of-function mutants in their ability to induce the Hh response *in cis*. This activation does not require processing of the Shh pro-protein, as a majority of the tested holoprosencephaly mutants in the large a-helix do not give rise to ShhN_Chol_ (Figure 5F). This suggests that at least some cases of holoprosencephaly are caused by the acquisition of cell autonomous, Ptch1/2-independent Hh signaling via the SHH pro-protein, while leaving non-autonomous signaling via the wild type allele intact.

**Figure 6.**
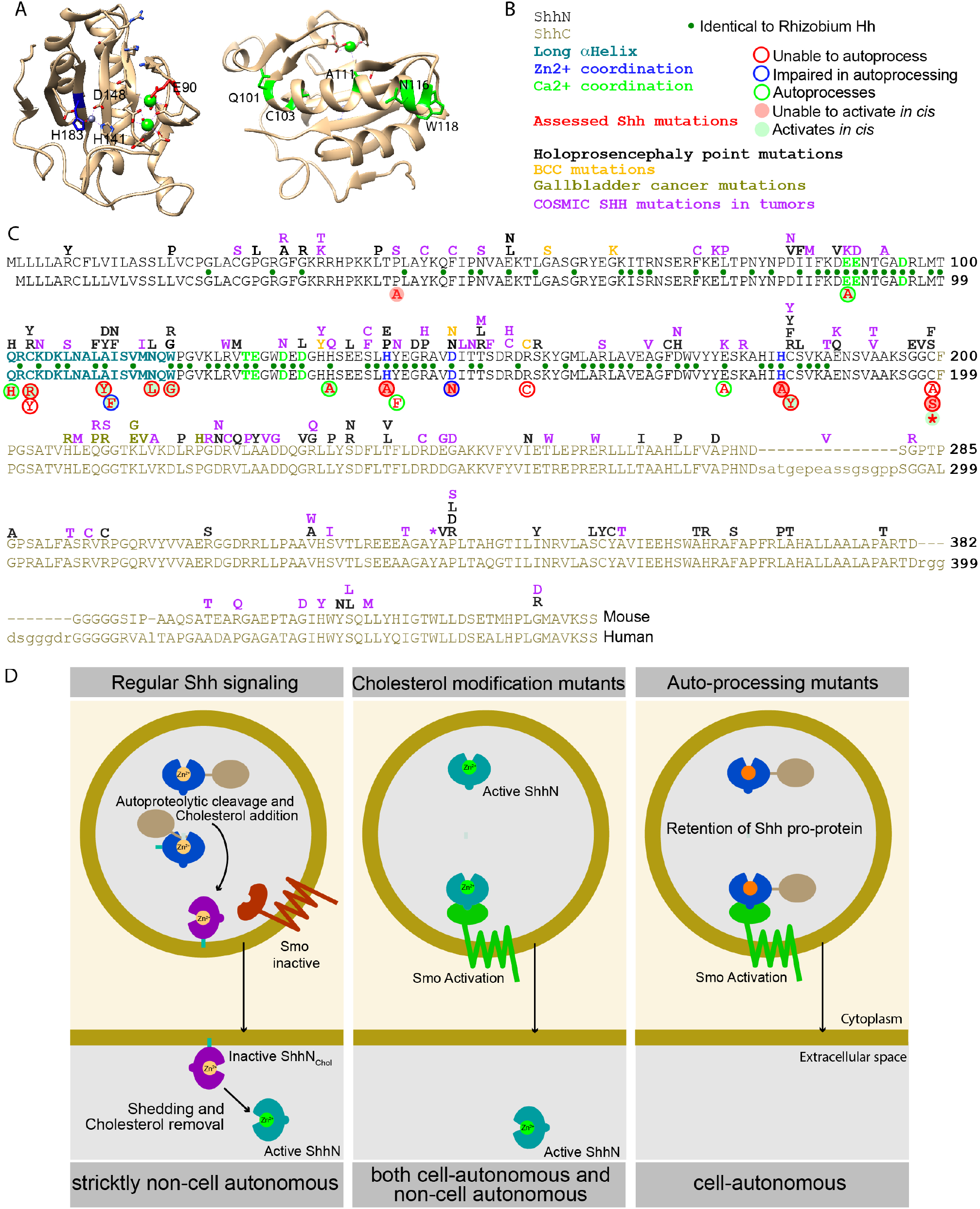
Overview and model. Representation of the ShhN crystal structure with salient residues indicated. On the left is a view into the Zn^2+^ (grey)/Ca^2+^ (green) coordination domains, on the right is the opposite view with the large α-helix in front, with mutated residues indicated in green. (B, C) Lineup of the mouse (top) and human Shh. All tested mutants are indicated below the lineup, with their activity and ability to process indicated. Above the line are Shh point mutations described for holoprosencephaly, BCCs, Gallbladder Cancers, and those curated in the COSMIC database. Identical residues found in *Mesorhizobium* Hh are indicated with green dots between the lines. (D) During regular Hh signaling, the potentially active pro-protein autocleaves to yield the inactive Shh_Chol_ form. Further processing mediates activation and release, thus allowing signaling *in trans*. Mutants that do not receive the cholesterol modification always exist in an active form, thus mediating both cell-autonomous and non-cell autonomous signaling. Repressed autoprocessing of the Shh proprotein results in perdurance of this active form, thus causing cell-autonomous activation of the Hh response.

## Discussion

### Ptch1/2-independent activation of Smo by Shh

Our observations challenge the canonical model that Smo activation is solely mediated by the Shh-induced release of Ptch1 inhibition. Despite the ample evidence that Ptch1 is an efficient inhibitor of Smo (Goodrich et al., 1997), we find that: 1) the loss of *Ptch1/2* does not inevitably result in maximal Smo activation, and 2) several forms of Shh can activate the migrational and transcriptional Hh responses in the absence of Ptch1/2 function. Nevertheless, Ptch1 function is required in those same cells to transcriptionally respond to ShhN provided extracellularly (Roberts et al., 2016). A central role of Ptch1 function is to regulate Smo entry into the primary cilium (Rohatgi et al., 2007), which is required for the transcriptional Hh response (Goetz and Anderson, 2010) but dispensable for the chemotactic response (Bijlsma et al., 2012). Our observations are consistent with a model in which Shh can both bind to Ptch1/2 (Rohatgi et al., 2007), resulting in the localization of Smo to the primary cilium where it mediates the transcriptional response, and activate Smo independent of Ptch1/2 function, resulting in fast changes in cell shape that underlie chemotaxis. This model would require the function of Ptch1/2 to couple extracellular Shh to the Gli-mediated transcriptional response, something that might be circumvented by some of Shh mutants we described.

Whether cells lacking Ptch1 or Ptch2 retain sensitivity to Shh *in vivo* is unclear. However, some observations indicate that they might. The spectrum of tumors observed in *Ptch1* null and in *Ptch1/2* heterozygous mice (Y. Lee et al., 2006) are commonly found at sites where Shh is expressed; and Shh-induced tumors often arise at the same locations (Beachy et al., 2004). The positional overlap between tumors induced by the loss of Ptch function and Shh-induced tumors is compatible with our model that cells lacking Ptch1/2 function nevertheless retain some ability to respond to Shh. Our model is further supported by the observation that the establishment of left/right asymmetry in mouse embryos, determined by *Pitx2* expression, requires *Smo, Shh*, and *Ihh*, but not *Ptch1* (X. M. Zhang et al., 2001). *Ptch1^−/−^* mutant embryos establish normal asymmetric expression of *Pitx2*, indicating that this Hh/Smo-mediated symmetry-breaking event occurs independently of Ptch1. Furthermore, the upregulation of pathway activity observed in *Ptch1*^LacZ/LacZ^ mice coincides with a widespread expression of Shh in the developing neural tube, leaving open the possibility that this activation remains Shh-dependent.

The ability of Shh to activate Smo in a relatively direct manner is supported by the observations that ShhN activates the Hh pathway in cells lacking all known extracellular receptors and that forms of Shh unable to bind to these receptors activate the pathway in a Smo-dependent manner. Furthermore, we demonstrate that the CRD of Smo is required for both Shh-mediated transcriptional and migrational responses. The idea that the CRD of Smo is a target of Shh-mediated activation is supported by the previous observations that Smo-ΔCRD has a decreased sensitivity to Shh *in vitro* and *in vivo* (Aanstad et al., 2009; Myers et al., 2013; Nachtergaele et al., 2013). The regulation of Smo facilitated by Ptch1/2-mediated inhibition likely targets the heptahelical domain of Smo, a notion that has previously been suggested based on the action of many small molecule Smo inhibitors, such as Cyclopamine (J. K. Chen et al., 2002) and vismodegib (Byrne et al., 2016). This indicates that the mechanism by which Smo is inhibited and activated are not the same, consistent with the findings that: 1) Ptch1 can inhibit forms of Smo lacking the CRD (Myers et al., 2013), and 2) the binding of oxysterols and cholesterol to the CRD can activate Smo (Huang et al., 2016; Nachtergaele et al., 2013; 2012; Nedelcu et al., 2013).

### Shh autoproteolysis is not required for Smo activation

Our results demonstrate that a wide variety of Shh mutants, several previously characterized as “dead”, have the ability to activate Smo *in cis* when expressed in cells lacking the canonical receptors in vitro and in wild-type cells in vivo. ShhC199A, which remains unprocessed as a full length precursor poorly signals *in trans* (Pettigrew et al., 2014; Roelink et al., 1995), but can induce the Hh response *in cis*. Several Shh mutations cause persistence of the pro-proteins (C103Y/R, A111D/Y I112F/N, N116L, W118R/G, and C184Y) and these mutants can activate Hh response *in cis*. These results indicate that one important function of the unusual processing and modification of Shh is to prevent activation of the Hh response in cells that express the ligand, thus reinforcing strict non-cell autonomy that characterizes Shh signaling (García-Zaragoza et al., 2012; Shaw et al., 2009; Yauch et al., 2008).

The Zn^2+^ coordination site is involved in the binding to Hhip (Bosanac et al., 2009) 5E1 (Maun et al., 2010) and possibly Ptch1. The resemblance to the active site of Zn^2+^ peptidases was recognized early (Hall et al., 1995b), and this site was thought be not catalytically active, but instead was proposed to mediate binding to Ptch1 (Fuse et al., 1999). The Zn^2+^ coordination domain is required for autoproteolytic processing, consistent with the notion that (similar to Thermolysin), the Zn^2+^ triad is catalytically active. At low expression levels the Zn^2+^ triad mutants are poor inducers of the Hh response at best. However, several other mutants outside the triad prevent autoproteolytic processing, and gain the ability to induce the Hh response *in cis*. This indicates that the Zn^2+^ coordination domain, even in the pro-protein, appears to be involved in the activation of the Hh response in cells lacking Ptch1/2, and might thus be involved in the Ptch1/2-independent activation of Smo.

The ShhN domain, the mature signaling peptide that contains the Zn^2^coordination site, has highly conserved homologs in bacteria (Accession numbers in the Materials and Methods). Identitcal encompasses all critical amino acids involved in metal coordination, identical residues between Hh found in *Rhizobium* and mouse/human are indicated in Figure 6C. These predicted bacterial Hh proteins are members of a larger group of prokaryotic peptidases that are characterized by the Hedgehog/DD-transpeptidase motif (van Heijenoort, 2011). The signature transpeptidase domain is a central, five-stranded antiparallel β-sheet surrounded by α-helices, as found in ShhN. Members of this family include murein endopeptidase (penicillin resistant enzymes that modify bacterial cell walls) and peptidase M15 (bacterial D-alanyl-D-alanine carboxypeptidases (DD-transpeptidases), the target for penicillin). Like in Shh, in all DD-transpeptidases Zn^2+^ is coordinated by two Histidine residues and an Aspartic acid residue in a H-X(6)-D and H-X-H motif (H141 D148, H181, H183 for Shh). DD-transpeptidases use a catalytic mechanism in which the acyl-linked peptide-enzyme intermediate is resolved by a nucleophilic attack of the adjacent peptidoglycan chain, thus crosslinking the peptidoglycan chains in the bacterial cell wall. This mechanism is possibly related to resolving the reaction intermediate that exists during Shh autocleavage by cholesterol nucleophilic attack, resulting in ShhN_Chol_. It is thus possible that the Hh/DD-transpeptidase fold has evolved to prevent resolution of the peptide-enzyme intermediate by OH^−^, resulting in proteolysis.

This model is not consistent with the one proposed by the Beachy lab that requires C199 to allow a thiol-ester intermediate to be resolved by cholesterol (Porter et al., 1996). Our observation that C199S undergoes some processing (Figure 5E) cannot be explained by the Beachy model, but is consistent with a function of Shh as auto-transpeptidase that auto-processes.

### Shh mutants found in tumors and in individuals with holoprosencephaly

It is striking that several mutations found in SHH-induced Basal Cell Carcinomas (BCCs) (Couvé-Privat et al., 2004) (SHH-D147N and SHH-R155C) are defective in auto-processing (Figure 5 and not shown). It is unlikely that these mutations are inhibitory to Hh signaling and consistent with the idea that such mutants are gain-of-function as measured by the cell-autonomous activation of the Hh response. SHH Point mutations curated in the COSMIC database tend to be more prevalent in the SHHN domain, but are common in the SHHC domain right after the cleavage site. Asymmetric distribution indicates that at least some of the mutations are “drivers” in tumor formation. Several of the N-domain mutations involve residues that activate the cell autonomous response when mutated (E89K, H134Q, and C183Y). Several point mutations in gall bladder tumors are found in SHHC (Dixit et al., 2017), and at least one of these (Shh-K213E) fails to process. These findings are consistent with a model in which SHH mutations permit *in cis* activation of the Hh response, thereby contributing to tumor progression.

In humans, heterozygous *SHH* mutants causing single amino acid substitutions can cause holoprosencephaly (Hehr et al., 2010), although mice heterozygous for a null *Shh* allele are normal. It is thus possible that some of these point mutations are dominant-negatives or gain-of-function mutations. The analysis of point mutations that cause holoprosencephaly (Roessler et al., 2009) reveal an interesting pattern. Residues involved the Zn^2+^ coordination, and thus likely required for the peptidase activity, are targets for point mutations in holoprosencephaly (H140, and D147, human numbering), but also heterozygous mutations in the C-terminal domain of SHH can cause holoprosencephaly (Hehr et al., 2010). Mutations in the C-terminal domain likely interfere with autoprocessing of full length Shh and cholesterol addition to G197 of SHH. It is thus possible that either unprocessed SHH or SHH lacking the putative peptidase activity function as a dominant-negative forms of Shh as proposed by Singh *et al*. (Singh et al., 2009). Alternatively, a cell-autonomous activation of the Hh response (a gain-of-function mutation) in cells heterozygous for this mutation could contribute to holoprosencephaly. We think the latter explanation is more plausible given the gain-of-function phenotypes we find associated with *Shh* mutants that fail to process, or lack the cholesterol moiety.

### Model

Based on our results in cells lacking Ptch1/2 and those of others, we propose the following model to account for the strict non-cell autonomy of Shh signaling (Figure 6 D). Although the Shh pro-protein has the ability to activate the Hh response, this is prevented by the processing into the inactive ShhN_Chol_ form. This form is activated upon Disp1, Scube2, and metalloproteases to be released in an active form (Jakobs et al., 2014) that signals *in trans*. Mutations that prevent efficient processing of the Shh pro-protein, or prevent the addition of the cholesterol moiety allow cell autonomous activation of the Hh response. The activation of this response does not require, but likely is antagonized by Ptch1/2, It does, however, require the CRD of Smo, perhaps indicating that Smo is a receptor for Shh.

## Materials and Methods

### Materials

Vismodegib was a gift from Dr. Fred de Sauvage (Genentech). SAG was from EMD Biochemicals. Recombinant ShhN protein was from R&D Systems. Cell Tracker Green CMFDA was from Invitrogen.

### Electroporations

Hamburger-Hamilton (HH) stage 10 *Gallus gallus* embryos were electroporated caudally in the developing neural tube using standard procedures (Meyer and Roelink, 2003). Embryos were incubated for another 48 h following electroporation, dissected, fixed in 4% PFA, mounted in Tissue-Tek OCT Compound (Sakura) and sectioned.

### Embryoid Body differentiation

mESCs were neuralized and differentiated into embryoid bodies (NEBs) using established procedures (Wichterle et al., 2002). NEBs were harvested after 5 days in culture, fixed, and stained for Nkx2.2 and Isl1/2 (Kawakami et al., 1997). NEBs were mounted in Fluoromount-G (Southern Biotech) and imaged.

### Immunofluorescence

Antibodies for mouse Pax7 (1:10), Mnr2 (1:100), Nkx2.2 (1:10), Shh (5E1, 1:20) were from the Developmental Studies Hybridoma Bank. The Rabbit α-GFP (1:1000) antibody was from Invitrogen, and the Goat α-hOlig2 (1:100) antibody was from R&D Systems. The mouse α-acetylated tubulin (1:200) was from Sigma Aldrich. Alexa488 and Alexa568 secondary antibodies (1:1000) were from Invitrogen. Nuclei were stained with DAPI (Invitrogen).

### DNA Constructs

The *Gli-Luciferase* reporter and the Renilla control were gifts from Dr. H. Sasaki (Sasaki et al., 1997). *Ptch1* was a gift from Dr. Scott (Stanford University, CA, USA). *Ptch1-Δloop2* was a gift from Dr. Thomas Jessell (Columbia University, NY, USA). *Ptch1* channel mutants were previously described (Alfaro et al., 2014a). SmoΔCRD was a gift from J. Reiter (Aanstad et. al., 2009). SmoM2 was from Genentech (F. de Sauvage). The following mutations were created using Quikchange mutagenesis (Stratagene): *Shh-E90A, Shh-H183A, ShhN-E90A, ShhN-H183A, SmoΔCRD-CLD, SmoL112DW113Y*. Dhh and Ihh were gifts from Charles Emerson Jr. (University of Massachusetts Medical School, MA, USA). *DhhN* and *IhhN* were made by site directed mutagenesis of C199 (Dhh) and C203 (Ihh) to a stop codon. ShhC199A was previously described (Roelink et al., 1995). Gli3^PHS^ was previously described (Meyer et. al., 2003).

### Genome Editing

TALEN constructs targeting the first exon of mouse Cdo and Gas1 were designed and cloned into the pCTIG expression vectors containing IRES puromycin and IRES hygromycin selectable markers (Cermak et al., 2011). The following repeat variable domain sequences were generated: Cdo, 5’ TALEN: NN HD NI NG HD HD NI NN NI HD HD NG HD NN NN; 3’ TALEN: HD NI HD NI NI NN NI NI HD NI NG NI HD NI NN; Gas1, 5’ TALEN: NN NI NN NN NI HD NN HD HD HD NI NG NN HD HD; 3’ TALEN: NN NN NI NI NI NI NN NG NG NG NN NG HD HD NN NI. Two CRISPR constructs targeting a double strand break flanking the first exon of mouse Boc were cloned into *pSpCas9* vector with an IRES puromycin selectable marker (Ran et al., 2013). The Boc CRISPRs targeted the following forward genomic sequences (PAM sequences underlined): Upstream of first exon 5’<u>CCT</u>GTCCTCGCTGTTGGTCCCTA 3’; Downstream of first exon 5’CCCACAGACTCGCTGAAGAGCTC 3’. *Ptch1*^LacZ/LacZ^;*Ptch2^−/−^;Shh^−/−^* mouse embryonic stem cells (Roberts et al., 2016) were plated at a density of 1.0×10^6^ on 6-well plates and transfected with 6 genome editing plasmids the following day. One day after transfection, selective ES medium (100 μg/mL hygromycin and 0.5 μg/mL puromycin) was added for 4 days. Selective medium was removed and surviving mESC colonies were isolated, expanded and genotyped by sequence PCR products spanning TALEN and CRISPR-binding sites.

### Genotyping

PCR screening was performed on cell lysates using primers flanking the TALEN or CRISPR binding sites for the *Boc, Cdo*, and *Gas1* loci. *Boc*, (5’) CATCTAACAGCGTTGTCCAACAATG and (3’) CAAGGTGGTATTGTCCGGATC; *Cdo*, (5’) CACTTCAGTGTGATCTCCAG and (3’) CCTTGAACTCACAGAGATTCG; Gas1, (5’) ATGCCAGAGCTGCGAAGTGCTA and (3’) AGCGCCTGCCAGCAGATGAG. PCR products were sequenced. Samples with signals indicative of INDEL mutations were cloned into the pCR-BluntII vector using the Zero Blunt TOPO PCR cloning kit (Invitrogen) and sequenced to confirm allele sequences. A *Ptch1*^LacZ/LacZ^;*Ptch2^−/−^;Boc^−/−^;Cdo^−/−^;Gas1^−/−^;Shh^−/−^* mESC clone was identified harboring a 50 bp deletion in Cdo exon 1, a heteroallelic 480 bp insertion and a 200 bp deletion in Gas1 exon1 resulting in a premature stop codon in the reading frame, and a 450 bp deletion of Boc exon 1.

### Cell Culture

*Ptch1*^LacZ/LacZ^;*Ptch2^−/−^, *Ptch1*^LacZ/LacZ^;Ptch2^−/−^;Smo^−/−^, Ptch1^−/−^;Disp1^−/−^;Shh^−/−^*, and *Ptch1*^LacZ/LacZ^;*Ptch2^−/−^;Boc^−/−^;Cdo^−/−^;Gas1^−/−^;Shh^−/−^* fibroblasts were obtained by plating mESCs at a density of 8.0×10^5^ cells in 6-well plates and transfected with the *large T antigen* from the SV40 virus (Gökhan et al., 1998) in ES medium. Cells were then switched to DMEM (Invitrogen) supplemented with 10% fetal bovine serum (FBS) without LIF. *Ptch1*^+/LacZ^ and *Ptch1*^LacZ/LacZ^ fibroblasts (gifts from Dr. Scott) were cultured in DMEM supplemented with 10% FBS (Invitrogen) and maintained under standard conditions. Identity of these lines was confirmed by the presence of the LacZ recombination in the Ptch1 locus, the presence of 40 chromosomes per cell, and mouse-specific DNA sequences of the edited genes. mESC lines were maintained using standard conditions in dishes coated with gelatin, without feeder cells. Cells were routinely tested for Mycoplasm by Hoechst stain, and grown in the presence of tetracycline and gentamycin at regular intervals. Cultures with visible extranuclear staining, likely infected with Mycoplasma, were discarded. None of the cell lines used in this study is listed in the Database of Cross-Contaminated or Misidentified Cell Lines. Fibroblast-like lines derived from the mESCs were re-sequenced at the edited loci to confirm their identity.

### Transfection

Cells were transiently transfected for 24h at 80-90% confluency using Lipofectamine 2000 reagent (Invitrogen) according to the manufacturer’s protocol.

### Modified Boyden Chamber Migration Assay

Cell migration assays were performed as previously described (Bijlsma et al. 2007). Cells were labeled with 10 μM CellTracker Green (Invitrogen) in DMEM for one hour. The well compartments were set up with the specified chemoattractant (ShhN .75 μg/ML (resuspended in 0.1%BSA in PBS), 10% FCS, or no attractant (plus 0.1%BSA in PBS) and pre-warmed at 37°C. Cells were then detached with 5mM EDTA and resuspended in DMEM without phenol red and supplemented with 50mM HEPES. Cells were transferred into FluoroBlok Transwell inserts (BD Falcon) at 5.0×10^4^ cells per insert. GFP-spectrum fluorescence in the bottom compartment was measured every 2 min for 99 cycles (approximately 3 hours), after which background fluorescence (medium without cells) and a no-attractant control was subtracted from each time point. Starting points of migration were set to 0.

### Gli-Luciferase Assay

Fibroblasts were plated at a density of 3×10^4^ in 24 well plates and transfected with *Gli-Luciferase, CMV-Renilla* (control plasmid), and specified plasmids 24 hours after plating. Cells were grown to confluency and then switched to low serum medium (0.5% FBS) alone or with specified concentrations of Vismodegib. After 24 hours, cells were lysed and the luciferase activity in lysates was measured using the Dual Luciferase Reporter Assay System (Promega). Raw Luciferase values were normalized against Renilla values for each biological replicate to control against variation in transfection efficiency. Individual luciferase/renilla values were then normalized against the mock control average for each experiment.

### LacZ Assay

Fibroblasts were plated at a density of 3×10^4^ in 24 well plates and transfected with plasmids 24 hours after plating. Fibroblasts were grown to confluency and then switched to a low serum medium (0.5% FBS) alone or with specified concentrations of Vismodegib or SAG. After 24 hours, cells were lysed and lysates were analyzed using the Galacto-Light™ chemiluminescence kit (Applied Biosciences) for level of LacZ expression. Raw chemiluminescence values were normalized against total protein for each biological replicate. Protein concentration was determined with a Bradford assay using the Bio-Rad Protein Assay Dye Reagent.

### Western Blots

*Ptch1*^LacZ/LacZ^;*Ptch1^−/−^* cells were transfected with Shh mutants as indicated. 48 hours after transfection, *Ptch1*^LacZ/LacZ^;*Ptch2^−/−^* cells were rinsed with PBS and lysed with RIPA buffer (150 mM NaCl, 50 mM Tris-HCl, 1% Igepal, 0.5% Sodium Deoxycholate, and protease inhibitors) for 30 min on ice. Protein lysate was cleared by centrifugation at 13,000g for 30 min at 4 °C. 20 μg of each sample was run on a 12% SDS-PAGE gel and transferred to a 0.45 micron nitrocellulose membrane. Membranes were blocked with 5% milk in Tris-buffered saline with 0.1% Tween-20 (TBS-T) and incubated with a rabbit polyclonal anti-Shh antibody (H160; Santa Cruz Biotechnology) at 1:250. A goat anti-rabbit HRP-conjugated secondary antibody (BioRad) was used at 1:10000.

### NCBI Reference Sequence Mesorhizobium HhN

WP_029077494.1

## Acknowledgements

This work was supported by NIH grant 1R01GM117090 to HR. Vismodegib was a gift from Dr. de Sauvage (Genentech), *Dhh* and *Ihh* were a gift from Dr. Charles P. Emerson III (U. Mass. Med. School). Dr. A. Alfaro cloned the Shh binding mutants (E90A, H183A) and made the initial observation that they activate the Hh pathway *in vivo*. We thank Dr. M. Barro for her help with Western blotting, Dr. M. F. Bijlsma (AMC Amsterdam) for his help with the migration assays, and Dr. B. Roberts (Allen Institute for Cell Science) for his comments and advice. We also thank Fatma Ozguc for help with genome editing of the *Ptch1*^LacZ/LacZ^;*Ptch2^−/−^;Boc^−/−^;Cdo^−/−^;Gas1^−/−^;Shh^−/−^* mESC line and Michelle Boisvert for her help with generating Shh mutants, and Carina Jägers and Wei Lu for commenting on the manuscript.

## Author contributions

CC performed nearly all experiments. HR and CC designed the experiments and wrote the manuscript.

